# OXA-1207, a new OXA-48-type carbapenem-hydrolysing class D beta-lactamase from *Escherichia coli*

**DOI:** 10.1101/2025.07.31.667721

**Authors:** Julian Sommer, Marie Welter, Felix F. Krause, Stephan Göttig

## Abstract

We present the characterisation of OXA-1207, a novel, plasmid-encoded OXA-48-like carbapenemase identified in two unrelated *Escherichia coli* (sequence type (ST) ST405 and ST4405) isolates from hospital patients in Germany. OXA-1207 is closely related to OXA-181, differing by two amino acid substitutions (214G and 244W). The *bla*_OXA-1207_ gene was located on a self-transmissible IncFIC plasmids (pOXA-1207) featuring a conserved backbone and the novel ΔTn*6361* transposon structure, which derived from IncX3 plasmids carrying *bla*_OXA-181_. Notably, pOXA-1207 exhibited an exceptionally high conjugation frequency (mean 1.2 × 10^−1^) to *E. coli*, surpassing the typical rates reported for widely distributed OXA-48 plasmids. Analysis of public genomic datasets identified 40 *E. coli* isolates carrying *bla*_OXA-1207_, belonging to diverse high-risk clonal lineages (ST167, ST405 and ST410). The isolates originated from multiple continents, indicating global dissemination. These findings highlight OXA-1207 as an emerging and concerning resistance determinant with enhanced transmissibility and emphasise the need for continued molecular surveillance of OXA-48-like carbapenemases.

## 2. Introduction

Enterobacterales resistant to carbapenem antibiotics are on the rise and pose a global public health threat. Infections by carbapenem-resistant Enterobacterales are difficult to treat because of the very limited therapeutic options available. The primary cause of carbapenem resistance in Enterobacterales is the presence of carbapenemases, which are bacterial enzymes that hydrolyse carbapenems and most other beta-lactam antibiotics. In Europe, the Middle East and North Africa, the most prevalent carbapenemase is OXA-48, an Ambler class D beta-lactamase. The encoding gene *bla*_OXA-48_ is most frequently identified on a highly conserved 63.3 kb plasmid of the incompatibility group L (IncL), which plays a critical role in the dissemination of this enzyme due to its high transmission potential by horizontal gene transfer (HGT) and a low fitness burden (1). In recent years, a growing number of at least 65 different OXA-48 variants with high sequence identity to OXA-48 have been identified, commonly referred to as OXA-48-like carbapenemases (2). These enzymes differ by only limited amino acid changes but exhibit variable beta-lactam hydrolysis patterns. While the OXA-48 carbapenemase is the most prevalent variant, the growing number of OXA-48-like carbapenemases gain significance by rapid spread in certain global areas (3). The mode of dissemination of the most frequent OXA-48-like carbapenemases varies; the spread of OXA-48 has been connected to the high transmission rate of a conserved IncL 63 kb plasmid with low fitness burden in *E. coli* and *K. pneumoniae* and high conjugation frequencies (1). Other *bla*_OXA-48_-like are found on different plasmid types like *bla*_OXA-232_ on mobilizable ColE plasmids and *bla*_OXA-181_ on IncX3 plasmids (4,5). In contrast to that, *bla*_OXA-244_ is predominantly identified chromosomally encoded in high-risk *E. coli* ST38 clones, causing clonal outbreaks in European countries (6). The genetic environment of the *bla*_OXA-48_-like genes is characterised by various insertion sequences that form composite transposons, such as Tn*1999* (*bla*_OXA-48_) or Tn*2013* (*bla*_OXA-181_). Due to the low minimal inhibitory concentrations (MICs) against carbapenems of isolates producing OXA-48-like carbapenemases, these isolates can be hard to detect in routine laboratory screening methods, enforcing a silent spread (7). The spread of OXA-48-like variants is therefore based on different mechanisms, including horizontal gene transfer, the spread of successful clonal lineages and the difficult to detect antibiotic resistance phenotype (3).

The specific characteristics of the genetic environment and the genetic background of these genes influence their potential for dissemination in different niches and to new, potentially high-risk pathogens.

In this study, we identified and characterised the novel carbapenemase OXA-1207 in *E. coli*. We present an analysis of the characteristics of OXA-1207, including its antibiotic resistance phenotype, genetic background and horizontal gene transfer. We also examine the epidemiology of this carbapenemase using publicly available databases.

## 3. Material and Methods

### Bacterial isolates and antimicrobial susceptibility testing

*Enterobacterales* clinical isolates carrying the plasmid-borne OXA-48-like carbapenemases OXA-48, OXA-484 and OXA-1207 were recovered from patients of the University Hospital Frankfurt, Germany (Table 1). All isolates were phenotypically characterized and tested for *bla*_OXA-48-like_ by PCR and Sanger sequencing as previously described (8). For detection of carbapenemases, the immunochromatographic CARBA 5 lateral flow test (NG-Biotech, Guipry, France) was used (9). Antimicrobial susceptibility was determined using antibiotic gradient strips and (Liofilchem, Roseto degli Abruzzi, Italy), broth micro dilution for temocillin, ertapenem, meropenem, imipenem and colistin and agar dilution for fosfomycin as recommended by EUCAST. MICs were interpreted according to EUCAST guidelines v15.0.

**Table 1:**
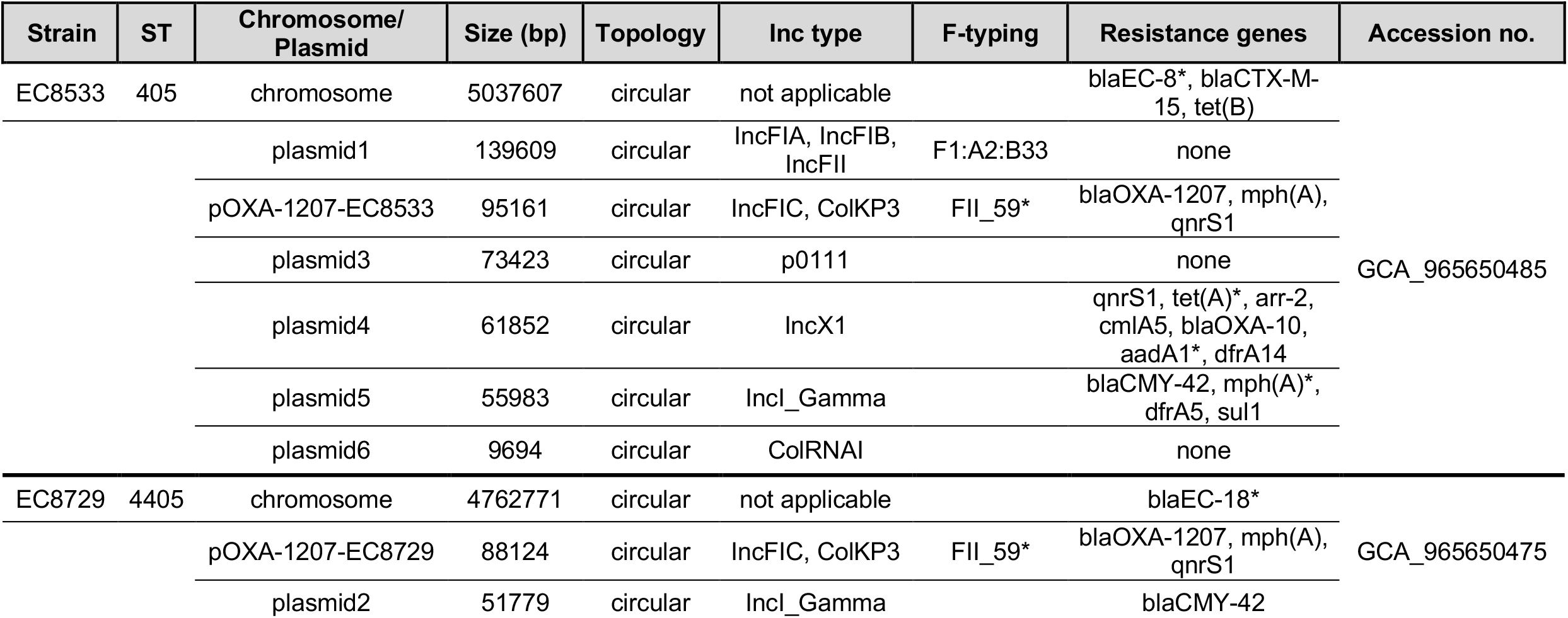
Molecular characteristics of the clinical isolates EC8533 and EC8729 encoding *bla*_OXA-1207_.

### Cloning of *bla*OXA-48-like

The open reading frames of *bla*_OXA-1207_ and *bla*_OXA-48_, including their native promoters, were amplified using the primers OXA-TOPO-F1 (AGTGTTGCTCTGTGGATAAC), preOXA-48B (CACACAAATACGCGCTAACC) and cloned into the expression vector pCR-Blunt II-TOPO (Invitrogen, Darmstadt, Germany) as described before (10,11). The resulting vectors pTOPO_OXA-1207 and pTOPO_OXA-48 were used to transform electrocompetent *E. coli* J53 and *K. quasipneumoniae* subsp. *quasipneumoniae* PRZ (12).

### Horizontal gene transfer of *bla*OXA-48-like harboring plasmids

Transconjugation of *bla*_OXA-48-like_ harboring plasmids was conducted by liquid mating as previously described (13). Briefly, a mixture of donors and recipients in brain heart infusion broth were incubated over night at 37°C and subsequently plated on chromogenic agar plates containing 100 mg/L sodium azide and 20 mg/L amoxicillin-clavulanic acid to select for transconjugants (Tc). Clinical *E. coli* isolates encoding *bla*_OXA-48_ or *bla*_OXA-1207_ were employed as donors and sodium azide-resistant *E. coli* J53 and *K. quasipneumoniae* subsp. *quasipneumoniae* PRZ as recipients (1). Presence of OXA-48-like carbapenemases in Tc was verified by disk diffusion antibiotic testing and the CARBA 5 lateral flow test. Randomly picked *E. coli* J53 transconjugants were analysed using disk diffusion testing and long-read sequencing. Transconjugation frequency was determined by dividing the numbers of Tc colonies by the number of acceptor colonies.

### Whole genome sequencing and bioinformatic analysis

Whole genome sequencing was carried out for all clinical isolates using short-read technology (MiSeq or NovaSeq platform, Illumina, San Diego, USA) and long-read technology (GridION platform, Oxford Nanopore Technologies, Oxford, UK). DNA was extracted from isolates using the DNeasy UltraClean Microbial Kit (Qiagen, Hilden, Germany). For Illumina sequencing, a v3 reagent kit was applied generating 150 bp paired-end reads. Library preparation for Nanopore sequencing was done using the SQK-NBD114.24 or SQK-NBD114.96 ligation sequencing kit. Sequencing was performed on a GridION or P2 Promethion sequencer utilizing a R10.4.1 flow cells. Raw signal data was base called and demultiplexed using the super accuracy base calling model of the dorado basecaller software v0.9.4. Raw data was filtered using trimmomatic for short reads and NanoFilt for long reads resulting in datasets of reads with an average genome coverage of at least 100-fold for short-reads and at least 50-fold for long reads (14,15). *De novo* hybrid assembly was conducted using autocycler version 0.3.0 (16). The obtained assemblies were annotated using bakta version 1.11 (17). Sequence types were determined using the software mlst v2.19.0 (18). ABRicate v1.0.1 was applied using the databases PlasmidFinder and NCBI AMRFinderPlus for identification of plasmid incompatibility groups and antibiotic resistance genes respectively, using thresholds of 100% gene coverage and ≥ 98% nucleotide sequence identity (19). ISfinder and mobile element finder was employed for annotation of insertion sequences (20,21). Plasmid sequences were aligned using MAFFT 7.490 and visualized using Geneious prime 2025.0.3 (22). Plasmid sequences were clustered against the PLSDB database using the screen approach via the PLSDB webpage interface and the 30 most closely related, circular plasmids were downloaded for further analysis of size, antibiotic resistance genes and Inc type (23). These plasmids were clustered with the two pOXA-1207 plasmids from this study using the pling software with default parameters and cluster networks were visualised using cytoscape using the edge-weighted spring embedded layout (24,25).

### Statistical analysis

For the comparison of transconjugation frequencies, continuous variables were assessed by Mann-Whitney *U* test. A *P*-value of <0.05 was considered significant.

### Ethics Statement

All bacterial strains were isolated as part of routine microbiological diagnostics and stored in an anonymized database. According to the ethics committee of the Hospital of Johann Wolfgang Goethe-University, Frankfurt am Main, no informed consent or ethical approval of the study is necessary.

## 4. Results

### Identification of clinical isolates harbouring *bla*OXA-1207

The novel carbapenemase OXA-1207 was identified in two distinct *E. coli* isolates (EC8533 and EC8729) from two different patients at the University Hospital Frankfurt in Germany (Table1). The isolates were identified due to reduced inhibition zone to ertapenem using disc diffusion during routine diagnostics screening for multi-resistant bacteria. Interestingly, the Vitek® 2 automated system for antibiotic susceptibility testing issued a warning regarding a potential carbapenemase in isolate EC8729, but not in isolate EC8533, based on the detected MICs. Both isolates were identified in a rectal swab. The patients both had a history of recent travel to India prior to hospital admission, but the isolation of the bacterial strains was 6 months apart and no association could be made between the two patients in the hospital.

The isolates exhibited resistance against most beta-lactam antibiotics, including ertapenem, piperacillin-tazobactam, third-generation cephalosporins, aztreonam, and ceftolozan-tazobactam, but were susceptible to meropenem, imipenem, ceftazidime-avibactam and cefiderocol (Table 2). The isolate EC8533 was resistant to most tested non-beta-lactam antibiotics, including ciprofloxacin, trimethoprim-sulfamethoxazole, and the aminoglycosides amikacin and tobramycin. The isolate EC8729 presented a less resistant phenotype with susceptibility against trimethoprim-sulfamethoxazole and all tested aminoglycosides gentamicin, amikacin and tobramycin (Table 2).

**Table 2:** MICs of the clinical isolates producing OXA-1207, transconjugants (Tc) and transformants (Tf) of the *E. coli* J53 recipient harbouring plasmids or the expression vector pTOPO encoding either *bla*_OXA-1207_ or *bla*_OXA-48_.

| beta-lactam | EC8533 | EC8729 | J53 | Tc-8533<br>J53 | Tc-8729<br>J53 | Tc pOXA-48<br>J53 | TF-pOXA-1207<br>J53 | TF-pOXA-48<br>J53 |
| --- | --- | --- | --- | --- | --- | --- | --- | --- |
| Amoxicillin +<br>Clavulanic acid | ≥256 | ≥256 | 8 | ≥256 | ≥256 | ≥256 | ≥256 | ≥256 |
| Ampicillin +<br>Sulbactam | ≥256 | ≥256 | 8 | ≥256 | ≥256 | ≥256 | ≥256 | ≥256 |
| Piperacillin +<br>Tazobactam | ≥256 | ≥256 | 2 | ≥256 | 64 | 64 | ≥256 | ≥256 |
| Temocillin | ≥1024 | ≥1024 | 4 | 512 | 512 | 256 | 1024 | ≥1024 |
| Cefuroxime | ≥256 | ≥256 | 4 | 8 | 8 | 8 | 8 | 8 |
| Cefotaxime | ≥256 | ≥256 | 0.12 | 0.25 | 0.25 | 0.25 | 0.25 | 0.25 |
| Ceftazidime | ≥256 | ≥256 | 0.25 | 0.25 | 0.25 | 0.12 | 0.25 | 0.25 |
| Ertapenem | 4 | 4 | 0.002 | 0.125 | 0.12 | 0.25 | 0.5 | 1 |
| Imipenem | 1 | 0.5 | 0.12 | 0.25 | 0.5 | 1 | 0.5 | 1 |
| Meropenem | 2 | 0.5 | 0.01 | 0.06 | 0.06 | 0.1 | 0.12 | 0.25 |
| Imipenem +<br>Relebactam | 0.5 | 0.5 |  |  |  |  |  |  |
| Ceftazidime +<br>Avibactam | 4 | 2 |  |  |  |  |  |  |
| Cefiderocol | 2 | 2 |  |  |  |  |  |  |
| Gentamicin | 1 | 1 |  |  |  |  |  |  |
| Tobramycin | 2 | 1 |  |  |  |  |  |  |
| Amikacin | 2 | 2 |  |  |  |  |  |  |
| Trimethoprim-Sulfamethoxazole | ≥32 | 0.125 |  |  |  |  |  |  |
| Ciprofloxacin | ≥32 | ≥32 |  |  |  |  |  |  |
| Levofloxacin | ≥32 | ≥32 |  |  |  |  |  |  |
| Colistin | 1 | 0.5 |  |  |  |  |  |  |

Using the CARBA 5 lateral flow test, production of an OXA-48-like enzyme was identified in both clinical isolates and the *bla*_OXA-48_-like open reading frames were amplified by PCR with specific primer for *bla*_OXA-48_-like and subsequent Sanger-sequenced. Sequence comparison of the sequences with the *bla*_OXA-48_-like gene sequences in the NCBI’s pathogen detection reference gene catalogue identified the *bla*_OXA-1207_ gene (NG_203409.1).

### Genome analysis of *E. coli* EC8533 and EC8729

Both *E. coli* bacterial strains carrying *bla*_OXA-1207_ were whole genome sequenced using a hybrid long-read and short-read sequencing approach. Assemblies of the two strains revealed circular chromosomes, six circular plasmids in strain EC8533 and two circular plasmids in strain EC8729 (Table 1). The complete nucleotide sequence assemblies of the isolates were deposited publicly in ENA linked to the study number PRJEB94527. Multi locus sequence typing identified two different sequence types (ST), ST405 for EC8533 and ST4405 for EC8729. Antibiotic resistance gene detection identified multiple beta-lactamase genes in the strain EC8533; the carbapenemase gene *bla*_OXA-1207_, the extended spectrum cephalosporinase genes *bla*_CTX-M-15_ and *bla*_CMY-42_ and the narrow spectrum beta-lactamases *bla*_OXA-10_ and *bla*_EC-8_. Additional antibiotic resistance genes included genes conferring resistance to fluoroquinolones (*qnrS1*), macrolides (*mph(A)*), tetracyclines (*tet(B)*), aminoglycosides (*aadA1*), rifampicin (*arr-2*), chloramphenicol (*cmlA5*), trimethoprim (*dfrA5, dfrA14*) and sulphonamides (*sul1*) (Table 1). In contrast, the isolate EC8729 encoded less antibiotic resistance genes; *bla*_OXA-1207_, the cephalosporinase *bla*_CMY-42_, the narrow spectrum beta-lactamase *bla*_EC-18_ and genes conferring resistance against macrolides (*mph(A)*) and fluroquinolones (*qnrS1*) (Table 1).

### Sequence analysis of pOXA-1207

*Bla*_OXA-1207_ was encoded on circular plasmids of 95,161 bp (pOXA-1207-EC8533, strain EC8533) and 88,124 bp (pOXA-1207-EC8729, strain EC8729) with 92.6% sequence identity (Figure 1). These plasmids differ by an in-frame deletion of 9 bp in *traD*, four silent single nucleotide variants (SNV) in *traG* and one SNV each in insertion sequence (IS) IS26 and a hypothetical protein in the smaller plasmid pOXA-1207-EC8729. Additionally, the plasmid pOXA-1207-EC8533 had three regions of 2,342 bp, 2,343 bp and 2,343 bp with 99% sequence identity, encoding the group II intron reverse transcriptase/maturase *ItrA*, which was not present in pOXA-1207-EC8729. Molecular typing of the two plasmids using the plasmidfinder database revealed two relaxase genes (*rep*), one of the IncFIC plasmid group and an additional fragment of 229 bp of the original 280 bp *rep* from ColKP3 plasmids. The subtyping of the IncF alleles of these plasmids using the pMLST software identified the IncFII allele 59 with 99.3% sequence identity.

**Figure 1.**
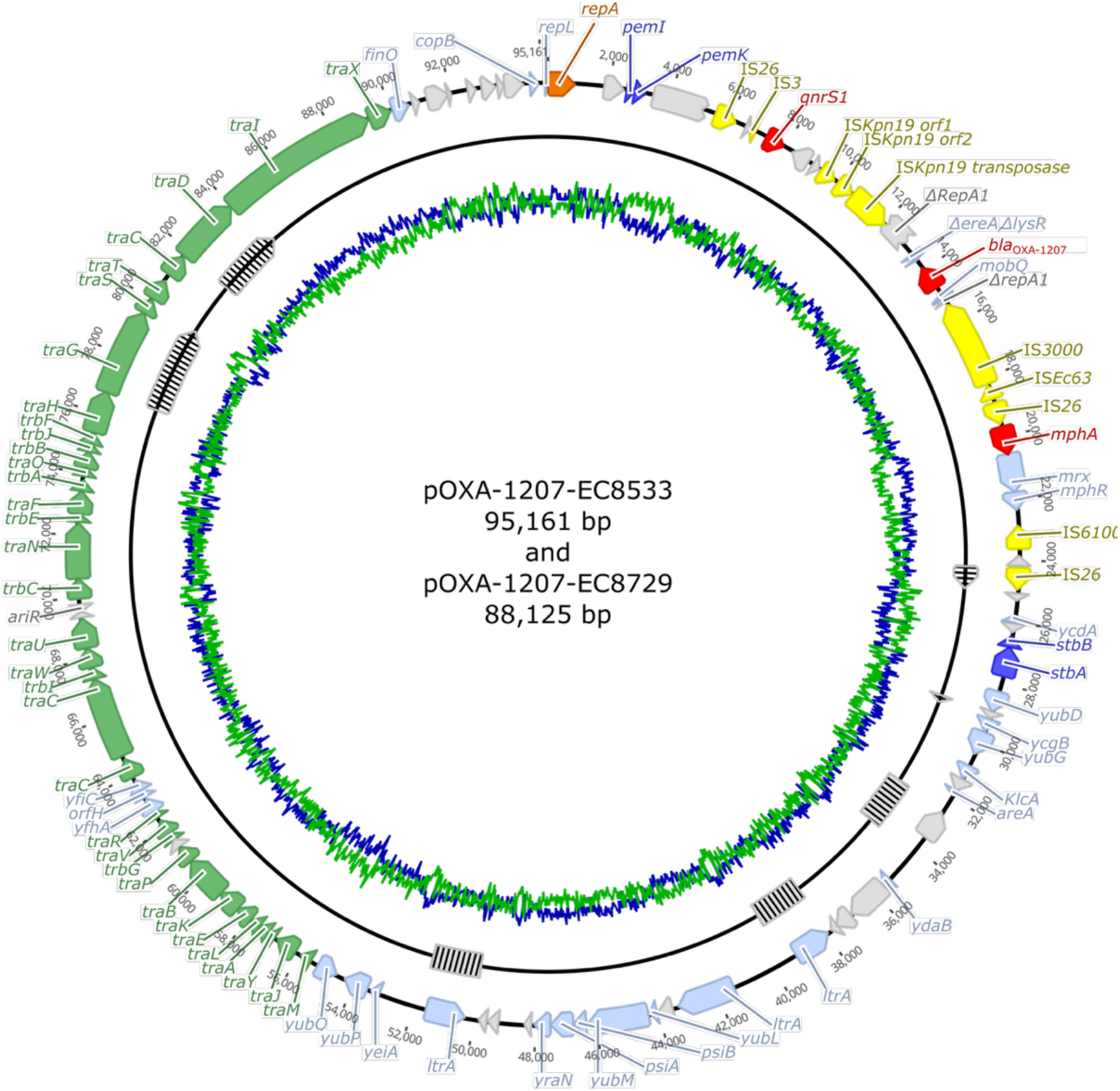
Plasmid map of pOXA-1207 from clinical isolate *E. coli* EC8533 (outer ring) with antibiotic resistance genes (red), insertion elements (IS, yellow), replication initiation protein A (orange), toxin-antitoxin systems (dark blue), disrupted genes and hypothetical proteins (grey), miscellaneous genes (light blue) and the *tra* locus of transfer-related genes (green). The differences between pOXA-1207-EC8729 and pOXA-1207-EC8533 are shown in the inner ring and are represented by striped boxes (deletions) and striped arrows (open reading frames with SNPs or small deletions). GC/AT-content: green/blue line.

The genetic environment of *bla*_OXA-1207_ on both plasmids was analysed using the ISfinder and MGE cluster software, identifying the resistant genes being embedded in a truncated Tn*6361* transposon structure of 12,785 bp length. Sequence comparison of this new ΔTn*6361* with the original Tn*6361* transposon from *bla*_OXA-181_ encoding plasmid (KM660724) showed a truncation of 2,671 bp on the upstream gene flanking region, one SNP in the IS26 and IS3000 open reading frames each and two SNPs in the *bla*_OXA-1207_ gene, compared to the *bla*_OXA-181_ gene in Tn*6361*. Upstream of the truncated ΔTn*6361*, a IS26-flanked *mph(A)-mpx-mphR* macrolide resistance gene cluster was identified (Figure 2). The *bla*_OXA-181_ harbouring Tn*6361* originates from the fusion of a Tn*2013* (ColKP3 plasmid, JN205800) and Tn*6292* from an IncN plasmid (pIMP-Z1058, KU051709) (Figure 2) (26).

**Figure 2.**
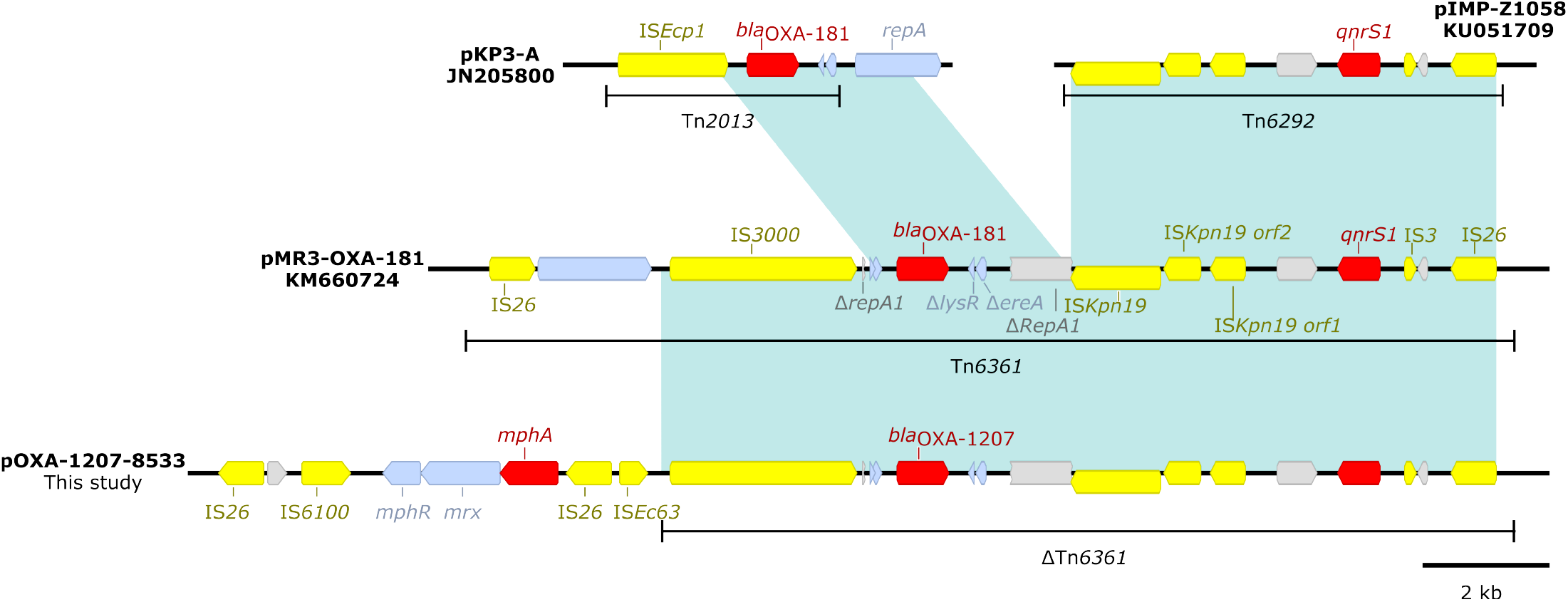
Genetic environment of natural plasmids of *bla*_OXA-1207_ and *bla*_OXA-181_. IS elements (yellow), fragmented genes (grey), antibiotic resistance genes (red) and other plasmid genes (light blue). Shaded regions share >99% of nucleotide sequence identity.

### OXA-1207 Plasmid clustering

To search for plasmids with high sequence similarity to the newly described pOXA-1207 plasmids, we used the clustering function of the plasmid database PLSDB and analysed the phylogenetic network of the two pOXA-1207 and the 30, circular plasmids from the database. The size of these plasmids ranged from 3,239 bp to 92,687 bp with a mean of 53,559 bp. Using the pling software to identify plasmid cluster networks in the identified plasmids and the two plasmids from this study, 29 of these were clustered together in a IncF-like cluster of plasmids, including the two plasmids from this study. These 29 plasmids included thirteen plasmids with no beta-lactamase gene present, ten plasmids with the narrow spectrum beta-lactamase *bla*_TEM-1_, two plasmids with *bla*_OXA-181_, the two plasmids from this study with *bla*_OXA-1207_, one plasmid with *bla*_SHV-12_ and one plasmid encoding the carbapenemase *bla*_NDM-5_. The three plasmids outside the IncF-like cluster included two plasmids without beta-lacatmase gene and one plasmid encoding *bla*_NDM-1_ (Figure 3).

**Figure 3.**
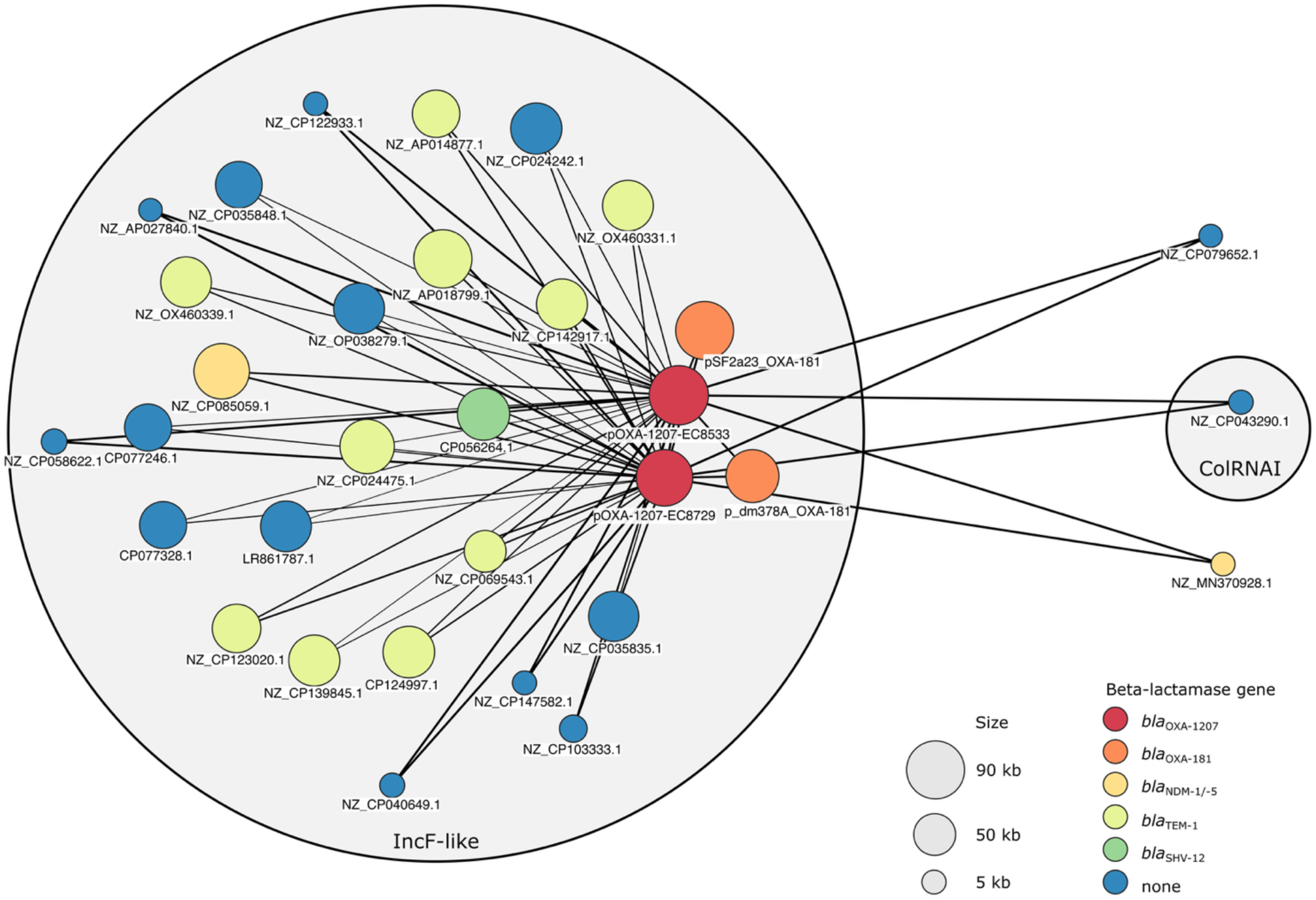
Plasmid cluster network of 32 plasmids, including the pOXA-1207 plasmids from this study and 30 most closely related plasmids from the PLSDB database. Clustering was performed using the plig software algorithm. The size of the circles encodes the plasmid size. The colour encodes beta-lactamase genes present on the plasmids. Gray large circles include plasmids of different Inc-types.

The two plasmids p_dm378A_OXA-181 (79,270 bp) and pSF2a23_OXA-181 (91,955 bp) encoded OXA-181 and had the highest mash identity of > 0.995 to pOXA-1207-EC8533 and originated from *E. coli* and *Shigella flexneri*, respectively. The two plasmids share the plasmid backbone of the pOXA-1207 plasmids with high sequence similarity and the ΔTn*6361* transposon structure including the *bla*_OXA-181_ gene; only 20 SNPs difference between pOX-1207-EC8533 and p_dm378A_OXA-181 in an IS*26* transposase, an IS*3000* transposase and the *bla*_OXA-1207_ / *bla*_OXA-181_ genes and 5 SNPs difference between pOXA-1207-EC8533 and the pSF2a23_OXA-181 in an IS*Kpn19* open reading frame, a hypothetical protein and the *bla*_OXA-1207_ / *bla*_OXA-181_ genes. The transposon structures flanking this ΔTn*6361* transposon, however, differ between these plasmids. On the plasmid p_dm378A_OXA-181, the IS*26*-flanked *mph(A)-mpx-mphR* gene cluster was not found. In plasmid pSF2a23_OXA-181, the IS*26*-flanked *mph(A)-mpx-mphR* gene cluster was located downstream of the *bla*_OXA-1207_ and *qnrS1* genes and an additional IS*26*-flanked transposon structure including the *emr(B)* gene coding for a multidrug efflux pump was present.

### Epidemiology of bacterial isolates producing OXA-1207 and plasmid support

Using the NCBI Pathogen Detection website, we queried for bacterial isolates carrying *bla*_OXA-1207_ in April 2025, identifying 39 isolates encoding *bla*_OXA-1207_ in total. We downloaded the reads or assemblies of the isolates when available, supplemented with additional metadata. These isolates included 38 *E. coli* isolates and one *Citrobacter amalonaticus* isolate; all but one environmental isolate were received from clinical samples. The earliest collection was in 2020 (n=1 isolate), all other isolates were collected in 2023 (n=12), 2024 (n=25) or 2025 (n=1). Most of the isolates were collected in the United States of America (n=27), five in Singapore, two in Vietnam, two in the Netherlands and one each in Canada, the United Kingdom and France (Figure 4). Phylogenetic analysis using a core genome SNP approach of all *E. coli* isolates and the two newly described *E. coli* isolates from this study revealed a heterogenic collection of isolates with thirteen different STs and one new ST. Of these, however, 23 (56%) isolates belonged to the four different high-risk STs ST167 (n=4), ST361 (n=4), ST405 (n=6) and ST410 (n=9). Analysis of the core genome single nucleotide polymorphisms (SNPs) of the isolates revealed potential cluster of clonal strains of three or more strains with 25 SNPs or less difference; one cluster of four isolates of ST410 with 8 to 23 SNPs difference including isolates from the USA and Vietnam, a second cluster of four isolates of ST361 with 0 to 2 SNPs from Singapore and a third cluster of four isolates of ST405 with 9 to 25 SNPs difference between isolates from the USA and the Netherlands (Figure 5).

**Figure 4.**
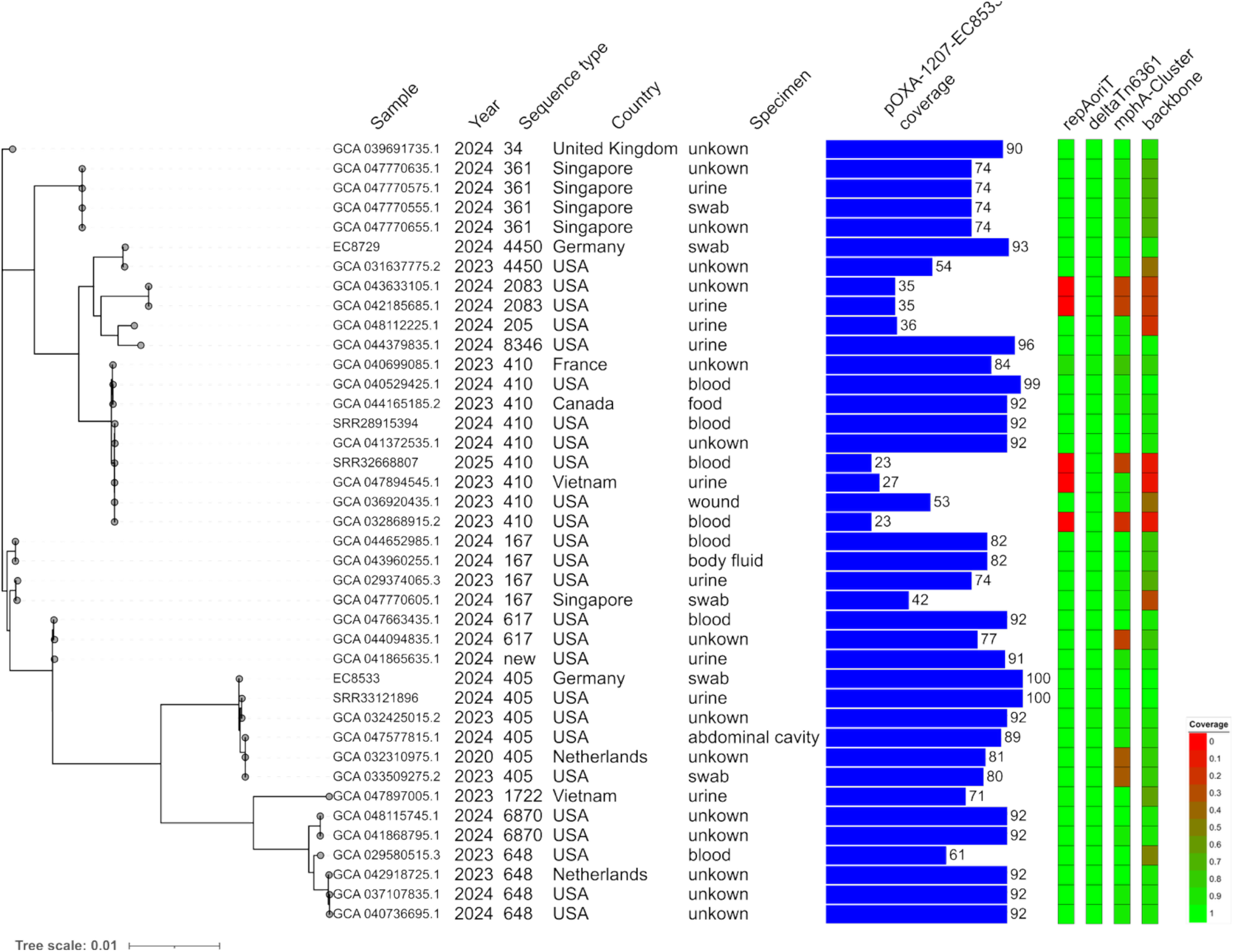
Phylogeny of *E. coli* strains producing OXA-1207 from this study (n=2) and additional isolates from the NCBI pathogen detection database (n=38). For each isolate, the year of isolation, the STs, country of collection and specimen is shown. The blue bars represent the coverage of the dataset of pOXA-1207-EC8533. The heatmap represent coverage of four regions of the plasmid pOXA-1207-EC8533: the *repAoriT* region (bases 1-1077), the ΔTn*6361* region including *bla*_OXA-1207_ (bases 5438-18222), the *mphA*-cluster (bases 18821-24923) and the plasmid backbone region (bases 25049-96161 and bases 1-5190).

**Figure 5.**
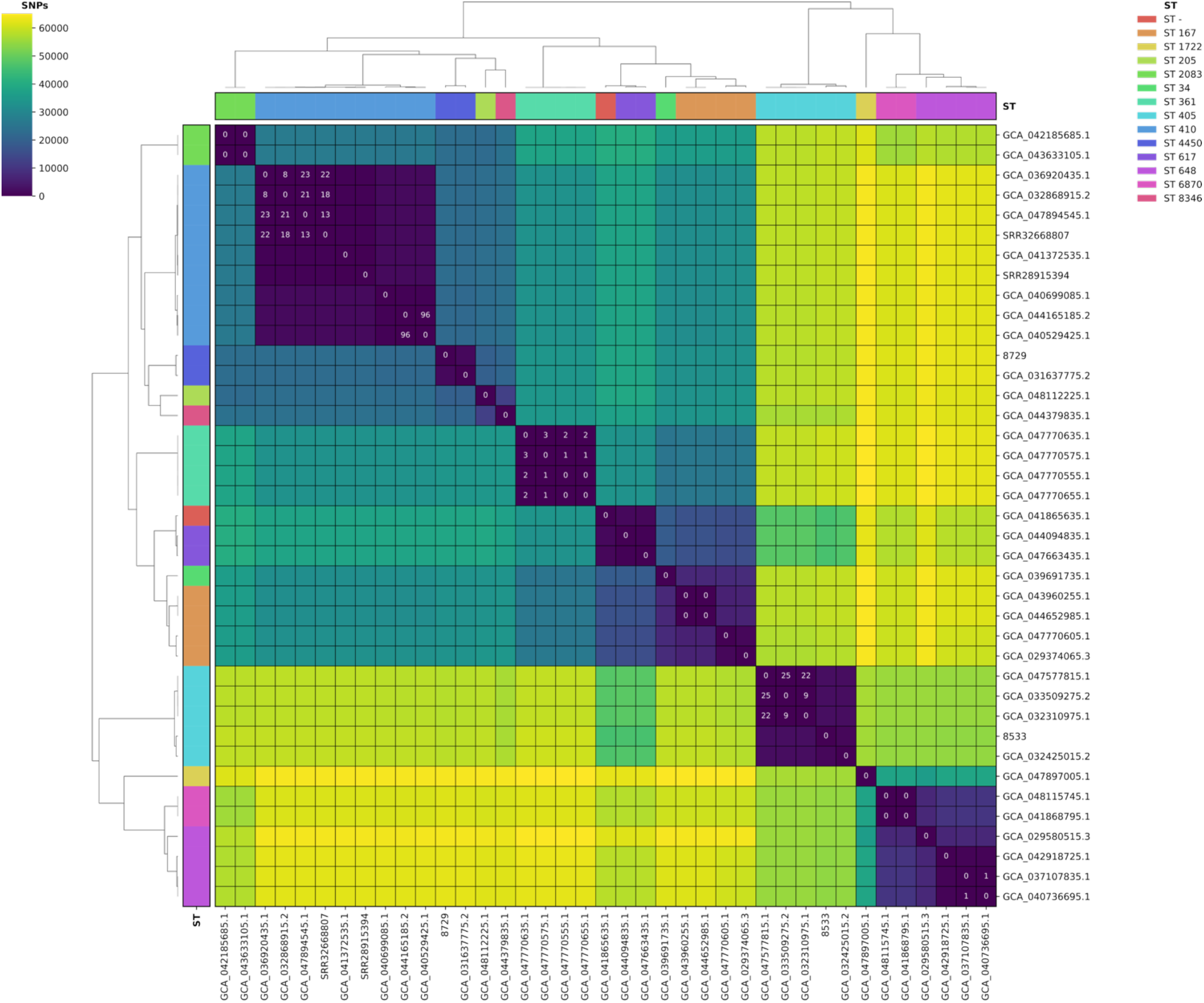
NP matrix of the core genome of 40 *E. coli* isolates producing OXA-1207. For each isolate, the number of SNP differences with all other isolates was calculated and used to cluster the isolates according to the Euclidean distance. The colour represents the number of SNPs separating the isolates. Values of ≤25 SNPs, indicating high similarity, are shown. The colour bar shows the sequence types of the isolates.

To analyse the genetic background of *bla*_OXA-1207_ in these datasets, we mapped the reads of the 39 datasets to the sequence of pOXA-1207-EC8533 and calculated the coverage of the complete plasmid. Additionally, we analysed the coverage in four defined regions of this plasmid, the sequence including the IncF replication initiation protein *repA* and the origin of replication *oriT* (bases 1-1077), the ΔTn*6361* including *bla*_OXA-1207_ (bases 5438-18222), the *mphA*-cluster (bases 18821-24923) and the plasmid backbone region (bases 25049-96161 and bases 1-5190). This data revealed a mean coverage of 74.6%, ranging from 23.1 to 100% for all read datasets; only seven of the 39 isolates had less than 50% coverage of the plasmid (Figure 4). Only five datasets had no coverage of the *repA*-*oriT* region suggesting no presence of a plasmid of the IncFIC type in these datasets. All datasets showed high coverage of the ΔTn*6361* region. Only two of the 39 datasets had coverage of less than 100% of this region, but both still showed coverage of more than 95% of this transposon structure. Thirty-two datasets mapped the *mph(A)*-cluster with a mean coverage of 94%, whereas seven datasets had coverage of less than 50% in this region. The backbone region of the pOXA-1207-EC8533 plasmid was covered to an average extent of 69% across all datasets, with only ten datasets covering less than 50% of the plasmid sequence. The five datasets lacking the IncFIC *repA-oriT* region showed coverage of the plasmid sequence of less than 25% (Figure 4).

### Horizontal gene transfer and plasmid stability of pOXA-1207

The transmission of pOXA-1207 from EC8533 and EC8729 to *E. coli* was analysed by *in vitro* liquid mating assays, using the sodium-azide resistant bacterial strain *E. coli* J53 as recipient. Transmission of pOXA-1207 to *E. coli* J53 was highly efficient with conjugation frequencies of 5.2×10^−1^ to 8.9×10^−1^ (Figure 6). The conjugation frequencies of the broadly disseminated 63.3 kb IncL plasmid encoding *bla*_OXA-48_ (pOXA-48) from two different *E. coli* donor strains (EC2700, EC5035) to *E. coli* J53 were significantly lower; EC2700 had a conjugation frequency of 2.8×10^−4^ to 2.0×10^−3^ and EC5035 of 8.9×10^−4^ to 1.3×10^−3^ (p< 0.005, Figure 6).

**Figure 6.**
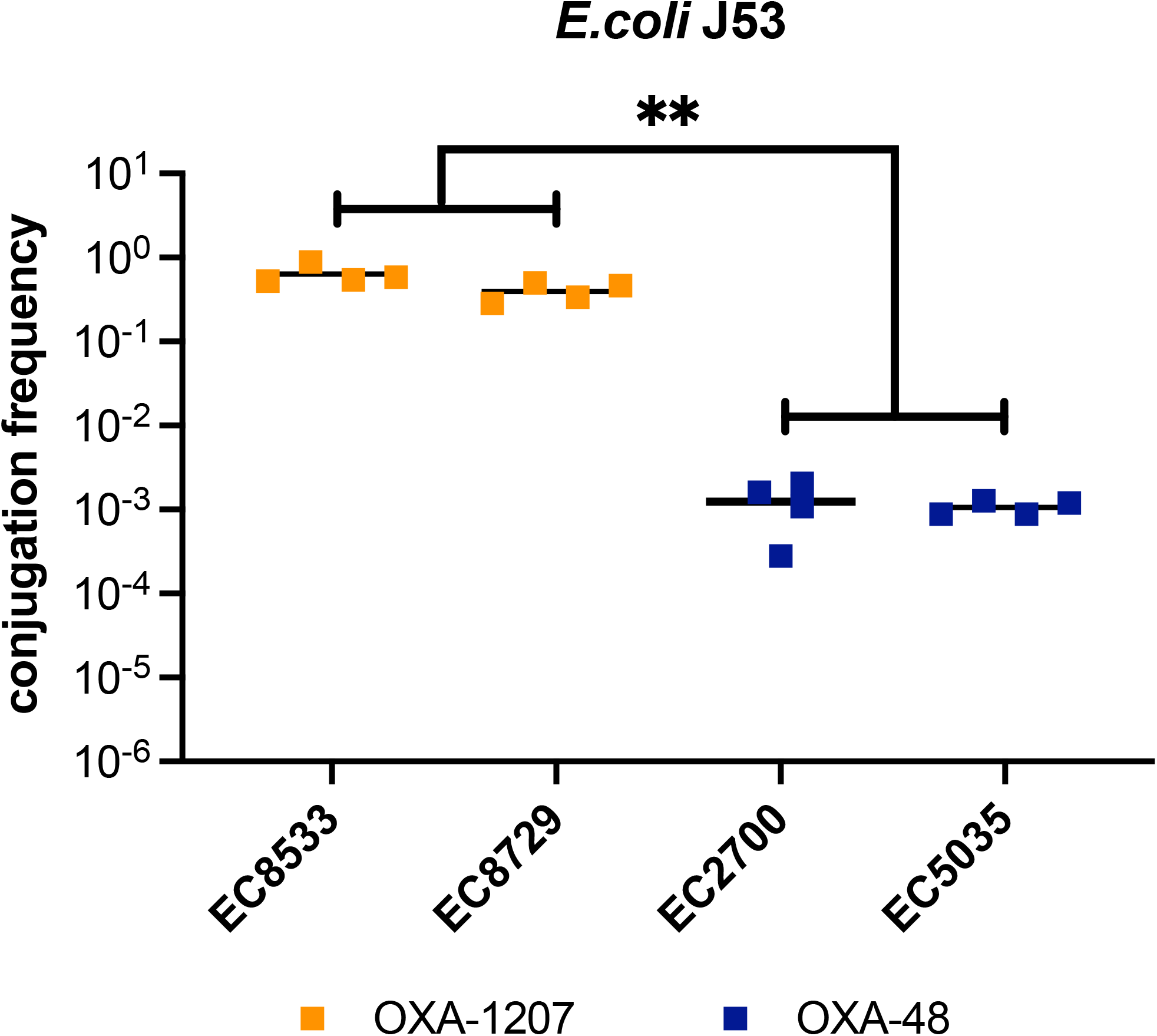
Conjugation frequencies of plasmids encoding *bla*_OXA-1207_ or *bla*_OXA-48_ from *E. coli* clinical isolates to the recipient *E. coli* J53. Each dot represents the frequency of a single conjugation experiment, whereas the bars indicate the mean of all experiments. **P < 0.005 (Mann-Whitney U test).

### Beta-lactam resistance mediated by OXA-1207

The hydrolytic activity of *bla*_OXA-1207_ was analysed determining the MICs of *E. coli* J53 transconjugants harbouring the plasmid pOXA-1207-EC8533, pOXA-1207-EC8729, or pOXA-48 from donor EC2700. In addition, the open reading frame of *bla*_OXA-1207_ and *bla*_OXA-48_ was amplified, cloned into the pTOPO expression vector and transformed into the recipient *E. coli* J53.

Analysis of the MICs of transconjugants compared to the MICs of the recipient without plasmids revealed an increase of the MICs to ampicillin, piperacillin ± tazobactam and temocillin of more than two MIC dilution steps, but no such significant increase of the MICs to cefuroxime, cefotaxime and ceftazidime (Table 2). The MICs of the carbapenems ertapenem, imipenem and meropenem increased in transconjugants producing OXA-48 or OXA-1207 by two or more MIC steps, except for imipenem in transconjugants harbouring pOXA-1207-EC8533.The MICs for carbapenems, however, were below the EUCAST cutoff for resistance for all carbapenems. Analysis of the transformants harbouring the pTOPO vector expressing *bla*_OXA-1207_ or *bla*_OXA-48_ presented comparable MICs, with increase of the MICs for carbapenemase by more than two dilution steps, with slightly higher MICs in transformants expressing *bla*_OXA-48_ compared to transformants expressing *bla*_OXA-1207_ (Table 2).

## 5. Discussion

In this study, we report the characterization of a novel OXA-48-like carbapenemase, OXA-1207, which was identified on an IncF plasmid type in two *E. coli* clinical isolates. Compared to the classical OXA-48-encoding IncL plasmids, the IncF plasmid harbouring *bla*_OXA-1207_ exhibits an unusually high conjugation frequency to *E. coli* recipients. Through *in silico* screening of publicly available databases, we identified 39 additional isolates encoding *bla*_OXA-1207_, originating from different continents and predominantly detected from 2023 onwards. Notably, 34 of the 39 isolates carried highly similar IncFIC plasmids, compared to those in the two isolates in this study. All but one of these isolates were from *E. coli*, including multiple isolates from high-risk clonal lineages, associated with the nosocomial spread of carbapenemases and human infections.

The two *E. coli* isolates producing OXA-1207 identified in this study exhibited the common antibiotic-resistant phenotypes of OXA-48-like-producing isolates, displaying resistance to only ertapenem but not meropenem and imipenem (Table 2). Laboratories that rely on the fully automated Vitek® 2 system for the detection of carbapenemase-producing Enterobacteria would have missed one of the two isolates. Therefore, the risk of silent spread of OXA-48-like producing Enterobacteria, especially with OXA-48-like variants with reduced β-lactamase hydrolysis, is high. The lateral flow test for detection of OXA-48-like carbapenemases, however, was positive for both isolates. Extended antibiotic susceptibility testing revealed cefazidime-avibactam and cefiderocol to be the only beta-lactam therapeutic options for the strains. One of the isolates belongs to the ST405, a high-risk sequence type carrying multiple virulence genes, that has been associated with severe infections and the spread of resistance to carbapenems and fluoroquinolones (27).

The two isolates share IncFIC-type plasmids with 93 % sequence similarity, including the plasmid backbone and the genetic environment of the three antibiotic-resistance genes, multiple insertion sequences and the newly described ΔTn*6361* transposon structure encoding *bla*_OXA-1207_ (Fig. 1). The two plasmids differ mainly in that the pOXA-1207-EC8533 plasmid contains three copies of group II intron maturases, which may have integrated into the plasmid itself (28). This highlights the similarity between the two plasmids. The ΔTn*6361* transposon structure is derived from the Tn*6361* transposon harbouring *bla*_OXA-181_, which has been identified on IncX3 plasmids. This plasmid type is mainly associated with the spread of OXA-181 (4). The identification of the two plasmids IncFIC-type plasmids pSF2a23_OXA-181 and p_dm378A_OXA-181 encoding *bla*_OXA-181_ with high sequence similarity to the OXA-1207 plasmids in this study, implies that the newly described ΔTn*6361* encoding originally *bla*_OXA-181_ has been integrated into IncFIC plasmids. Mutations in the open reading frame, accompanied by further modifications of the plasmid backbone, then led to the IncFIC plasmids encoding *bla*_OXA-1207_ (Fig. 2). Clustering of the pOXA-1207-EC8533 with plasmids from the PLSDB plasmid database revealed a cluster of 27 plasmids including the two previously mentioned *bla*_OXA-181_-encoding plasmids, which cluster closely with those from this study (Fig. 3). Interestingly, ten of these plasmids encode the beta-lactamase TEM, one of the most frequently identified beta-lactamase in *E. coli* (29). The spread of *bla*_TEM_ has been linked to highly transmissible IncF plasmids (30). The identification of the carbapenemase *bla*_OXA-1207_ on similar plasmids could indicate the potential for the efficient spread of these plasmids and the new carbapenemase OXA-1207 to different *E. coli* lineages. The detection of *bla*_OXA-181_ on very similar plasmids (pSF2a23_OXA-181) in a *Shigella flexneri* isolate suggests the possible spread of these IncFIC plasmids to additional species beside *E. coli*. The spread of plasmids, harbouring either *bla*_OXA-181_ or *bla*_OXA-1207_, to the obligate pathogen *Shigella* spp. would have a substantial impact on public health especially in low- and middle-income countries (31,32). Interestingly, the two IncFIC plasmids in this study carry the macrolide antibiotic resistance gene *mph(A)* in a IS26-*mph(A)*-*mrx*-*mphR(A)*-IS6100 mobile genetic element, which has been found in *Shigella* isolates from Bangladesh before (33). Therefore, the antibiotic resistance genes of the IncFIC plasmids could lead to extensively drug-resistant (XDR) *Shigella* spp. isolates via transmission, as they confer resistance to macrolide, fluroquinolone and beta-lactam antibiotics.

Phylogenetic analysis of the 39 isolates (38 *E. coli* and one *C. amalonaticus*) encoding *bla*_OXA-1207_ from the NCBI pathogen detection platform revealed that the isolates were collected from different regions of the world, mostly from 2023 onwards (Fig. 4). The absence of isolates collected before 2020 suggests a recent mutation event, most likely in the open reading frame of *bla*_OXA-484_, which differs from *bla*_OXA-1207_ by only one SNP. Interestingly, more than half of the isolates were collected in the United States of America, despite OXA-48-like carbapenemases accounting for only 10 % of total carbapenemases from Enterobacterales in the US (34). Additional isolates were also collected in south-east Asia, Europe and Canada, indicating the spread of OXA-1207 to different global regions, despite its first detection being relatively recent, in 2020, in the Netherlands (Fig. 4). Detailed phylogenetic analysis revealed three potential clusters of clonal isolates, which may indicate transmission or nosocomial outbreaks by clonal strains (Suppl. Figure 1). The twelve isolates from these clusters were identified as part of internationally disseminated, high-risk clonal lineages of *E. coli* (ST405, ST410 and ST361), which has been linked to the spread of antibiotic resistance and severe infections (35–37). The presence of OXA-1207 in these and other high-risk clonal lineages (ST167 and ST648) may increase the risk of this carbapenemase spreading in nosocomial environments (38,39). Most of the isolates, however, were not connected to a potential clonal cluster and belonged to 14 different STs, indicating a different mode of dissemination to that of clonal lineages spreading only (Fig. 4). Analysis of the plasmid sequences present revealed a high coverage of the IncFIC pOXA-1207 plasmids from this study for 34 (87%) of the isolates, suggesting the presence of a highly similar plasmid, potentially encoding *bla*_OXA-1207_ in most of the isolates (Fig. 4). In only five datasets was the presence of the IncFIC-defining *repA-oriT*-region not identified, suggesting the absence of this plasmid type in these isolates. The newly described ΔTn*6361* mobile genetic element, which encodes *bla*_OXA1207_, was found in all 39 isolates, including in the five in which the IncFIC plasmid type was not detected. This demonstrates a firm link between the spread of *bla*_OXA-1207_ and ΔTn*6361* and the potential capability of ΔTn*6361* to mobilise in new genetic context.

The spread of OXA-48 has been linked to its ability to efficiently conjugate the conserved IncL plasmid to a wide range of Enterobacterales without significant impact on the fitness or virulence of *E. coli* and *K. pneumoniae* host cells (1). High conjugation frequencies are also essential for the stable presence of antibiotic resistance plasmids in bacterial communities in the absence of antibiotic selection pressure (40). We identified conjugation frequencies to *E. coli* that were three orders of magnitude higher for IncFIC plasmids encoding *bla*_OXA-1207_ than for IncL plasmids encoding *bla*_OXA-48_ (Fig. 5). This finding could explain the rapid dissemination of the new carbapenemase OXA-1207 and its connection to IncFIC plasmids in a variety of different clonal *E. coli* lineages in a relatively short time. The presence of OXA-1207 in multiple high-risk clonal lineages could be an additional risk factor for increased spread in nosocomial environments and difficult to treat infections. These characteristics could indicate a high-risk plasmid-carbapenemase combination with substantial potential for further dissemination, which would exacerbate the global antimicrobial resistance crisis in Enterobacterales.

One limitation of our study is, that we did not perform enzyme kinetics of OXA-1207 to compare its hydrolysis activity with that of other OXA-48-like enzymes. Nevertheless, the increase of MICs to beta-lactams in transconjugants of the natural plasmids and transformants harbouring expression vectors encoding *bla*_OXA-1207_ showed comparable antimicrobial activity to that of OXA-48 (Table 2). Future studies, however, should analyse the therapeutic efficiency of carbapenems against strains expressing OXA-1207 in infection models *in vivo*.

## Literature

1. Hamprecht A, Sommer J, Willmann M, Brender C, Stelzer Y, Krause FF, et al. Pathogenicity of Clinical OXA-48 Isolates and Impact of the OXA-48 IncL Plasmid on Virulence and Bacterial Fitness. Front Microbiol. 2019 Nov 1;10:2509.

2. Naas T, Oueslati S, Bonnin RA, Dabos ML, Zavala A, Dortet L, et al. Beta-lactamase database (BLDB) – structure and function. J Enzyme Inhib Med Chem. 2017 Jan 1;32(1):917–9.

3. Peirano G, Pitout JDD. Rapidly spreading Enterobacterales with OXA-48-like carbapenemases. J Clin Microbiol. 2025 Jan 6;63(2):e01515–24.

4. Yu Z, Zhang Z, Shi L, Hua S, Luan T, Lin Q, et al. In silico characterization of IncX3 plasmids carrying blaOXA-181 in Enterobacterales. Front Cell Infect Microbiol [Internet]. 2022 Sept 8 [cited 2025 July 14];12. Available from: https://www.frontiersin.org/journals/cellular-and-infection-microbiology/articles/10.3389/fcimb.2022.988236/full

5. Potron A, Rondinaud E, Poirel L, Belmonte O, Boyer S, Camiade S, et al. Genetic and biochemical characterisation of OXA-232, a carbapenem-hydrolysing class D β-lactamase from Enterobacteriaceae. Int J Antimicrob Agents. 2013 Apr;41(4):325–9.

6. Kohlenberg A, Svartström O, Apfalter P, Hartl R, Bogaerts P, Huang TD, et al. Emergence of Escherichia coli ST131 carrying carbapenemase genes, European Union/European Economic Area, August 2012 to May 2024. Eurosurveillance. 2024 Nov 21;29(47):2400727.

7. Rima M, Emeraud C, Bonnin RA, Gonzalez C, Dortet L, Iorga BI, et al. Biochemical characterization of OXA-244, an emerging OXA-48 variant with reduced β-lactam hydrolytic activity. J Antimicrob Chemother. 2021 Aug 1;76(8):2024–8.

8. Göttig S, Gruber TM, Stecher B, Wichelhaus TA, Kempf VAJ. In Vivo Horizontal Gene Transfer of the Carbapenemase OXA-48 During a Nosocomial Outbreak. Clin Infect Dis. 2015;60(12):1808–15.

9. Boutal H, Vogel A, Bernabeu S, Devilliers K, Creton E, Cotellon G, et al. A multiplex lateral flow immunoassay for the rapid identification of NDM-, KPC-, IMP- and VIM-type and OXA-48-like carbapenemase-producing Enterobacteriaceae. J Antimicrob Chemother. 2018 Apr 1;73(4):909–15.

10. Potron A, Nordmann P, Lafeuille E, Al Maskari Z, Al Rashdi F, Poirel L. Characterization of OXA-181, a carbapenem-hydrolyzing class D β-lactamase from Klebsiella pneumoniae. Antimicrob Agents Chemother. 2011;55(10):4896–9.

11. Sommer J, Gerbracht KM, Krause FF, Wild F, Tietgen M, Riedel-Christ S, et al. OXA-484, an OXA-48-Type Carbapenem-Hydrolyzing Class D β-Lactamase From Escherichia coli. Front Microbiol. 2021 May 12;12(May):1–9.

12. Choi KH, Kumar A, Schweizer HP. A 10-min method for preparation of highly electrocompetent Pseudomonas aeruginosa cells: Application for DNA fragment transfer between chromosomes and plasmid transformation. J Microbiol Methods. 2006 Mar 1;64(3):391–7.

13. Gruber TM, Göttig S, Mark L, Christ S, Kempf VAJ, Wichelhaus TA, et al. Pathogenicity of pan-drug-resistant Serratia marcescens harbouring blaNDM-1. J Antimicrob Chemother. 2014;70(4):1026–30.

14. Bolger AM, Lohse M, Usadel B. Trimmomatic: A flexible trimmer for Illumina sequence data. Bioinformatics. 2014 Aug 1;30(15):2114–20.

15. De Coster W, D’Hert S, Schultz DT, Cruts M, Van Broeckhoven C. NanoPack: Visualizing and processing long-read sequencing data. Bioinformatics. 2018;34(15):2666–9.

16. Wick RR, Howden BP, Stinear TP. Autocycler: long-read consensus assembly for bacterial genomes [Internet]. bioRxiv; 2025 [cited 2025 June 18]. p. 2025.05.12.653612. Available from: https://www.biorxiv.org/content/10.1101/2025.05.12.653612v1

17. Schwengers O, Jelonek L, Dieckmann MA, Beyvers S, Blom J, Goesmann A. Bakta: rapid and standardized annotation of bacterial genomes via alignment-free sequence identification. Microb Genomics. 2021;7(11):000685.

18. Carattoli A, Zankari E, Garciá-Fernández A, Larsen MV, Lund O, Villa L, et al. In Silico detection and typing of plasmids using plasmidfinder and plasmid multilocus sequence typing. Antimicrob Agents Chemother. 2014;58(7):3895–903.

19. Feldgarden M, Brover V, Haft DH, Prasad AB, Slotta DJ, Tolstoy I, et al. Validating the AMRFINder tool and resistance gene database by using antimicrobial resistance genotype-phenotype correlations in a collection of isolates. Antimicrob Agents Chemother. 2019;63(11):e00483–19.

20. Siguier P, Perochon J, Lestrade L, Mahillon J, Chandler M. ISfinder: the reference centre for bacterial insertion sequences. Nucleic Acids Res. 2006 Jan 1;34:D32–6.

21. Johansson MHK, Bortolaia V, Tansirichaiya S, Aarestrup FM, Roberts AP, Petersen TN. Detection of mobile genetic elements associated with antibiotic resistance in Salmonella enterica using a newly developed web tool: MobileElementFinder. J Antimicrob Chemother. 2021 Jan 1;76(1):101–9.

22. Katoh K, Standley DM. MAFFT multiple sequence alignment software version 7: Improvements in performance and usability. Mol Biol Evol. 2013;30(4):772–80.

23. Molano LAG, Hirsch P, Hannig M, Müller R, Keller A. The PLSDB 2025 update: enhanced annotations and improved functionality for comprehensive plasmid research. Nucleic Acids Res. 2025 Jan 6;53(D1):D189–96.

24. Frolova D, Lima L, Roberts LW, Bohnenkämper L, Wittler R, Stoye J, et al. Applying rearrangement distances to enable plasmid epidemiology with pling. Microb Genomics. 2024;10(10):001300.

25. Shannon P, Markiel A, Ozier O, Baliga NS, Wang JT, Ramage D, et al. Cytoscape: A Software Environment for Integrated Models of Biomolecular Interaction Networks. Genome Res. 2003 Nov;13(11):2498–504.

26. Cuicapuza D, Alvarado L, Tocasca N, Aguilar D, Gómez-de-la-Torre JC, Salvatierra G, et al. First Report of OXA-181-Producing Enterobacterales Isolates in Latin America. Microbiol Spectr. 2023 June 15;11(3):e0458422.

27. Matsumura Y, Yamamoto M, Nagao M, Ito Y, Takakura S, Ichiyama S. Association of Fluoroquinolone Resistance, Virulence Genes, and IncF Plasmids with Extended-Spectrum-β-Lactamase-Producing Escherichia coli Sequence Type 131 (ST131) and ST405 Clonal Groups. Antimicrob Agents Chemother. 2013 Oct;57(10):4736–42.

28. Lambowitz AM, Zimmerly S. Group II Introns: Mobile Ribozymes that Invade DNA. Cold Spring Harb Perspect Biol. 2011 Aug 1;3(8):a003616–a003616.

29. Souza SSR, Piper KR, Ikhimiukor OO, Marcovici MM, Zac Soligno NI, Harmon AJ, et al. Variants of β-lactamase-encoding genes are disseminated by multiple genetically distinct lineages of bloodstream Escherichia coli. Commun Med. 2025 July 1;5(1):260.

30. Bradford PA. Extended-Spectrum β-Lactamases in the 21st Century: Characterization, Epidemiology, and Detection of This Important Resistance Threat. Clin Microbiol Rev. 2001 Oct;14(4):933–51.

31. Dhabaan G, Jamal H, Ouellette D, Alexander S, Arane K, Campigotto A, et al. Detection of OXA-181 Carbapenemase in Shigella flexneri. Emerg Infect Dis. 2024 May;30(5):1048–50.

32. Baker S, Scott TA. Antimicrobial-resistant Shigella: where do we go next? Nat Rev Microbiol. 2023;21(7):409–10.

33. Asad A, Jahan I, Munni MA, Begum R, Mukta MA, Saif K, et al. Multidrug-resistant conjugative plasmid carrying mphA confers increased antimicrobial resistance in Shigella. Sci Rep. 2024 Mar 23;14(1):6947.

34. Boyd SE, Holmes A, Peck R, Livermore DM, Hope W. OXA-48-Like β-Lactamases: Global Epidemiology, Treatment Options, and Development Pipeline. Antimicrob Agents Chemother. 2022 July 20;0(0):e00216–22.

35. Wang M, Zhang Z, Sun Z, Wang X, Zhu J, Jiang M, et al. The emergence of highly resistant and hypervirulent Escherichia coli ST405 clone in a tertiary hospital over 8 years. Emerg Microbes Infect. 14(1):2479048.

36. Ba X, Guo Y, Moran RA, Doughty EL, Liu B, Yao L, et al. Global emergence of a hypervirulent carbapenem-resistant Escherichia coli ST410 clone. Nat Commun. 2024 Jan 12;15(1):494.

37. Jousset AB, Bouabdallah L, Birer A, Rosinski-Chupin I, Mariet JF, Oueslati S, et al. Population Analysis of Escherichia coli Sequence Type 361 and Reduced Cefiderocol Susceptibility, France. Emerg Infect Dis. 2023 Sept;29(9):1877–81.

38. Johnson JR, Johnston BD, Gordon DM. Rapid and Specific Detection of the Escherichia coli Sequence Type 648 Complex within Phylogroup F. J Clin Microbiol. 2017 Mar 24;55(4):1116–21.

39. Garcia-Fernandez A, Villa L, Bibbolino G, Bressan A, Trancassini M, Pietropaolo V, et al. Novel Insights and Features of the NDM-5-Producing Escherichia coli Sequence Type 167 High-Risk Clone. mSphere. 2020 Apr 29;5(2):10.1128/msphere.00269-20.

40. Lopatkin AJ, Meredith HR, Srimani JK, Pfeiffer C, Durrett R, You L. Persistence and reversal of plasmid-mediated antibiotic resistance. Nat Commun. 2017 Nov 22;8:1689.

